# Homeodomain-interacting protein kinase (Hipk) is required for nervous system and muscle structure and function

**DOI:** 10.1101/719765

**Authors:** Simon Wang, Donald A.R. Sinclair, Hae-Yoon Kim, Stephen D. Kinsey, Byoungjoo Yoo, Kenneth Kin-Lam Wong, Charles Krieger, Nicholas Harden, Esther M. Verheyen

## Abstract

Homeodomain-interacting protein kinases (Hipk) have been previously associated with cell proliferation and cancer, however, their effects in the nervous system are less well understood. We have used *Drosophila melanogaster* to evaluate the effects of altered Hipk expression on the nervous system and muscle. Using genetic manipulation of Hipk expression we demonstrate that knockdown and over-expression of Hipk produces early adult lethality, partially due to the effects on the nervous system and muscle involvement. We find that optimal levels of Hipk are critical for the function of dopaminergic neurons and glial cells in the nervous system, as well as muscle. Furthermore, manipulation of Hipk affects the structure of the larval neuromuscular junction (NMJ) and increases motor neuron axonal branching. Hipk regulates the phosphorylation of the synapse-associated cytoskeletal protein Hts (adducin) and modulates the expression of two important protein kinases, Calcium-calmodulin protein kinase II (CaMKII) and Partitioning-defective 1 (PAR-1), all of which may alter neuromuscular function and influence lethality. Hipk also modifies the distribution of an important nuclear protein, TBPH, the fly orthologue of TAR DNA-binding protein 43 (TDP-43), which may have relevance for understanding motor neuron diseases.

## Introduction

Homeodomain-interacting protein kinase (Hipk) family members constitute a group of serine threonine kinases that phosphorylate many homeodomain transcription factors and influence cell proliferation and tissue growth (Blaquiere and Verheyen, 2017; Kim et al., 1998). In *Drosophila*, the single Hipk family member is essential for embryonic and larval survival and when over-expressed, acts as a potent growth regulator that can stimulate tumorigenesis and metastatic cell behavior through its actions on many signaling pathways including Wnt/Wingless, Hippo, Notch and JNK (Blaquiere et al., 2018; Chen and Verheyen, 2012; Huang et al., 2011; Kim et al., 1998; Lee et al., 2009a; Poon et al., 2012; Swarup and Verheyen, 2011). In *C. elegans*, the single Hipk ortholog, HPK-1, does not appear to be essential during development, but an *HPK-1* mutant significantly shortens adult lifespan (Berber et al., 2016). Vertebrates possess four Hipk orthologs that have some conserved and additional divergent functions (reviewed in Blaquiere and Verheyen, 2017). Numerous studies suggest that Hipk family members may also influence signaling pathways related to nervous system development and function. Flies with reduced dHipk possess small eyes with neuronal abnormalities (Blaquiere et al., 2014; Lee et al., 2009b). *Hipk1 Hipk2* double knockout mice are embryonic lethal and have underdeveloped retinas and defects in neural tube closure (Aikawa et al., 2006; Inoue et al., 2010; Isono et al., 2006). Mouse Hipk2 regulates the survival of sensory and sympathetic neurons, and midbrain dopamine neurons (Doxakis et al., 2004; Wiggins et al., 2004; Zhang et al., 2007a). In the embryonic midbrain, loss of mouse Hipk2 leads to a decrease in the number of neurons, resulting in Parkinson’s disease-like psychomotor abnormalities (Zhang et al., 2007a).

The relative level of expression of the *hipk* gene in the nervous system of larvae and adults is robust as described in Flybase (Thurmond et al., 2019). For these reasons, we explored the potential involvement of Hipk in the structure and function of the *Drosophila* nervous system. We show that down-regulation or over-expression of Hipk pan-neurally, as well as in dopaminergic neurons, glial cells and muscle cells is lethal. However, Hipk mis-expression in motor neurons, cholinergic or glutamatergic neurons has no obvious effects on viability. To investigate the potential causes of lethality from altered expression of Hipk, we focus on larvalneuromuscular junctions (NMJ) and muscle and find that Hipk regulates motor axon outgrowth and several synaptic proteins including Hts/adducin, Calcium/calmodulin-dependent protein kinase II (CaMKII) and PAR-1 (Partitioning-defective 1; a fly orthologue of microtubule affinity-regulating kinase). These studies demonstrate that Hipk regulates NMJ and muscle organization.

## Results

### Modulation of Hipk in adults leads to premature death

Since it has been shown that a *C. elegans HPK-1* mutant shortens adult lifespan (Berber et al., 2016), we decided to investigate the role of Hipk in adult *Drosophila* using the Gal4-UAS expression system in combination with a temperature sensitive *Gal80*^*ts*^ repressor to over-express and to knock-down Hipk after eclosion. We grew flies at 18°C until adulthood, and then shifted them to 29°C to inactivate the *Gal80*^*ts*^ repressor and to optimize Gal4 transcriptional activation of the UAS constructs. To rule out sex differences we used males in all adult survival analyses. For most studies, we utilized the 17090R-1 RNAi line from the National Institute of Genetics (NIG) to knock down *hipk* gene expression and will refer to this as *UAS-hipk-RNAi*, unless otherwise noted. We first assessed viability of adult males following *hipk* knockdown. The vast majority of *UAS-hipk-RNAi/Gal80*^*ts*^; *Tub-Gal4/*+ males died within 2-4 days after they were shifted from 18°C to 29°C (Fig. 1, red curves). With ubiquitous over-expression of Hipk using the HA-tagged *UAS-hipk*^3M^ transgene, (hereafter referred to as *UAS-hipk*), all *Gal80*^*ts*^*/*+; *UAS-hipk/Tub-Gal4* males died within approximately 7-10 days after they were shifted from 18°C to 29°C (Fig. 1, green curves). Typically, for both over-expression and knockdown experiments, there was a very small number of experimental flies that escaped early death. In both cases, prior to death, the activity of experimental males slowed progressively until they appeared immobile. In contrast, identically-treated internal control (Fig. 1, blue curves) and Oregon-R control (Fig. 1, grey curves) males survived considerably longer. These data indicate that both RNAi-induced knockdown and over-expression of Hipk dramatically affects survival and movement of males, suggesting the possibility that the flies may have impaired neural, and/or muscle function.

**Fig. 1:**
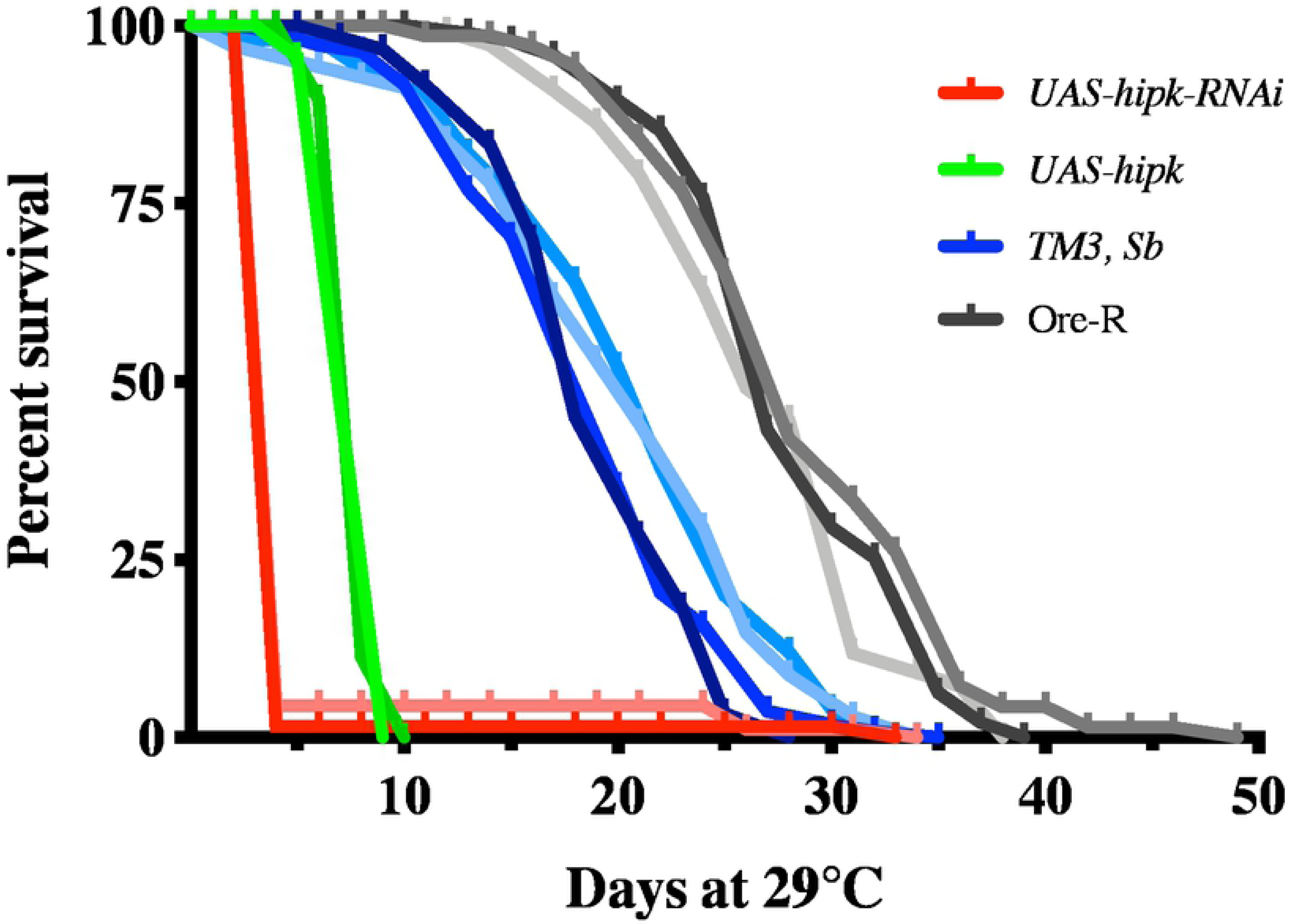
Ubiquitous modulation of Hipk affects adult life span. Adult males subjected to Hipk knockdown (*UAS-hipk*^*RNAi*^: NIG *UAS-hipk-RNAi*/*Gal80*^ts^; *Tub-Gal4*/+; light red and dark red curves; n= 89 and n= 65) died within 2 to 4 days at 29°C. Adult males subjected to Hipk over-expression (*UAS-hipk-RNAi*: 3M *UAS-hipk*/*Gal80*^ts^; *Tub-Gal4*/+; purple curves; n= 69 and n= 107) died within 7-10 days at 29°C. In contrast, Ore-R control males (n= 110; n= 96 and n= 69; light grey, dark grey and intermediate grey curves) died primarily within 25 to 40 days at 29°C, whereas internal control males (*TM3, Sb*: *UAS-hipk-RNAi*/*Gal80*^ts^; *TM3, Sb*/+ or *Gal80*^ts^/+; *UAS-hipk*/*TM3, Sb*; blue-purple curves; n= 69, n= 50, n= 109 and n= 60) died primarily within 21 to 25 days at 29°C.

### Modulation of Hipk during development affects specific cell types in the nervous system, as well as muscle cells

Based on these observations, we examined developmental effects of Hipk knockdown and over-expression at 29°C using several nervous system *Gal4* drivers and *Mef2-Gal4*, a muscle-specific driver (Tables 1, 2). Pan-neural knockdown of Hipk with *Appl-Gal4* resulted in complete lethality with a polyphasic lethal phase. To dissect this effect further we knocked down gene expression in subsets of neurons, namely dopaminergic, cholinergic, glutamatergic and motor neurons. As shown in Table 1, knockdown in dopaminergic neurons using *ple-Gal4* and *TH-Gal4* caused larval/pupal lethality, while knockdown in the other neurons did not affect viability. Knockdown using the pan-neural driver *elav-Gal4* resulted in rough eyes in adults (Fig. 2B). In addition, it is notable that Hipk knockdown in glial cells was also lethal, with a larval lethal phase. Finally, Hipk knockdown in muscle cells caused lethality at the pupal stage.

**Table 1:** Tests for viability of *hipk* knockdown in various nervous system components and muscles using the NIG *UAS-hlpk-RNAi*.

| *Gal4* Driver | Expression | % Progeny |  | Total Flies | Comments on *UAS-hipk-RNAi* |
| --- | --- | --- | --- | --- | --- |
|  |  | *UAS-hipk-RNAi* | Control (Balancer) |  |  |
| *Appl* | Pan-neural | 0 | 100 | 100 | Polyphasic LP, black structures in ventral termini of pupae |
| *GawB elav<sup>C155</sup>* | Pan-neural | 16.4 | 83.6 | 165 | Pupal LP; variably rough eyes with black patches |
| *ple* | Dopaminergic | 0 | 100 | 255 | Larval/pupal LP (undifferentiated) |
| *TH* | Dopaminergic | 4.3 | 95.7 | 115 | Larval/pupal LP (undifferentiated) |
| *ChAT-Gal4.7.4-19B* | Cholinergic | 54.2 | 45.8 | 96 | Viable |
| *ChAT-DD.Gal4.VP16* | Cholinergic | 51.1 | 48.9 | 278 | Viable |
| *GawB V Glut<sup>OK371</sup>* | Glutamatergic | 44.1 | 55.9 | 177 | Viable |
| *GawB D42* | Motor-neuron | 50.2 | 49.8 | 229 | Viable |
| *repo* | Glial cells | 0 | 100 | 155 | Larval LP |
| *Mef2* | Muscle | 0 | 100 | 157 | Pupal LP (undifferentiated) |

**Table 2:** Tests for viability of *hipk* over-expression in various nervous system components and muscles using the 3M *UAS-HA-hipk*.

| *Gal4* Driver | Expression | % Progeny |  | Total Flies | Comments on *UAS-hipk* |
| --- | --- | --- | --- | --- | --- |
|  |  | *UAS-HA-hipk* | Control (Balancer) |  |  |
| *Appl* | Pan-neural | 0.9 | 99.1 | 106 | Larval/pupal LP |
| *GawB elav<sup>C155</sup>* | Pan-neural | 26.9 | 73.1 | 67 | Rough eyes |
| *ple* | Dopaminergic | 0 | 100 | 225 | Polyphasic LP, some undifferentiated black pupae |
| *TH* | Dopaminergic | 0 | 100 | 158 | Larval/pupal LP (undifferentiated) |
| *ChAT-Gal4.7.4-19B* | Cholinergic | 55.5 | 44.5 | 164 | Viable |
| *ChAT-DD.Gal4.VP16* | Cholinergic | 51.1 | 48.9 | 278 | Viable |
| *GawB V Glut<sup>OK371</sup>* | Glutamatergic | 54.3 | 45.7 | 175 | Viable |
| *GawB D42* | Motor-neuron | 40.1 | 59.9 | 142 | Viable |
| *repo* | Glial cells | 0 | 100 | 174 | Larval LP |
| *Mef2* | Muscle | 0 | 100 | 156 | Larval/pupal LP (undifferentiated) |

**Fig. 2:**
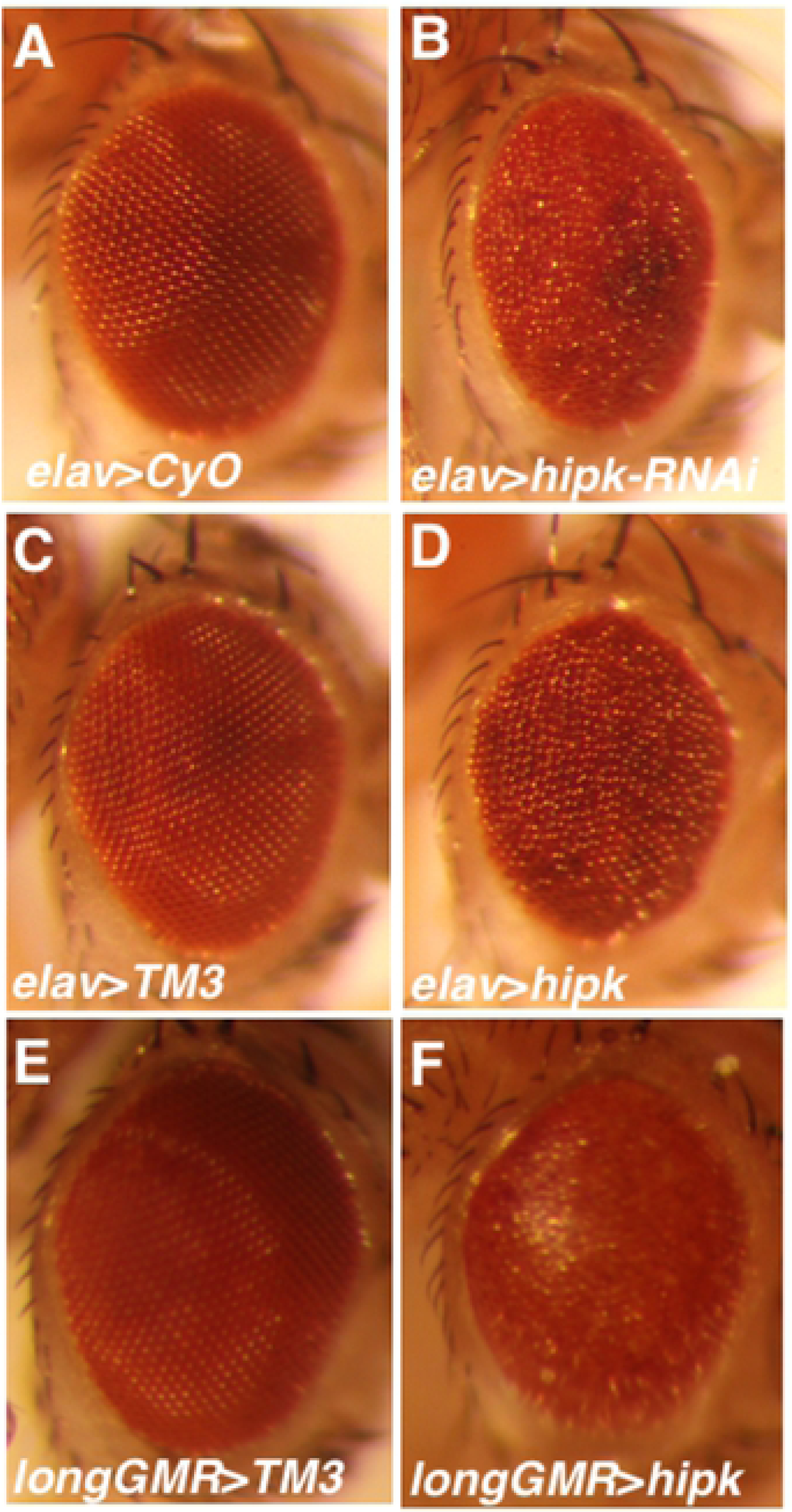
Disrupted eye patterning due to modulation of Hipk at 29°C. The eyes of internal control females (A: *elav*/+; *CyO*/+; C: *elav*/+; *TM3, Sb*/+; E: *longGMR-Gal4*/+; *TM3, Sb*/+) are normal. In contrast, the eyes of (B) *elav*/+; *UAS-hipk-RNAi*/+ and (D) *elav*/+; *UAS-hipk* (3M)/+ females are rough, whereas those of (F) *longGMR-Gal4*/+; *UAS-hipk*/+ females are smooth and have somewhat reduced pigmentation. No *UAS-hipk-RNAi*/*longGMR-Gal4* flies survived at 29°C or 25°C (data not shown).

To complement the knockdown experiments, we used the same *Gal4* driver strains to assess the effects of over-expression of Hipk during development using *UAS-hipk* (Table 2). We observed lethality at larval and pupal stages upon pan-neural expression using *Appl-Gal4*. In contrast, use of *elav-Gal4* resulted in rough eyes (Fig. 2D). Moreover, Hipk over-expression using *UAS-hipk* in combination with *longGMR-Gal4* generated flies with an abnormal glassy eye/reduced pigmentation phenotype (Fig. 2F). These studies highlight that both gain or loss of Hipk in neurons can affect eye formation and we have previously shown that Hipk is required during larval development for eye specification (Blaquiere et al., 2014). Furthermore, as was the case for Hipk knockdown, over-expression in dopaminergic neurons (larval lethal phase, LP), glial cells (larval LP) and muscle cells (larval/pupal LP) was lethal, but over-expression in cholinergic, glutamatergic or motor neurons was not.

Since Hipk knockdown in muscles and dopaminergic neurons during development was lethal, we carried out a second set of adult survival experiments to see whether such knockdown affected adult lifespan. For comparison, we included tests of ubiquitous over-expression of Hipk in adults. The data show that Hipk knockdown in muscle cells does indeed cause early adult death; nearly all of the *UAS-hipk-RNAi*/+; *Mef2-Gal4*/*Gal80*^*ts*^ males died within approximately 11-15 days at 29°C, in comparison to the control males (Fig. S1, purple versus black curves). However, Hipk knockdown in dopaminergic neurons using *TH-Gal4* males had lifespans similar to those of control males (Fig. S1, yellow versus black curves). Data for ubiquitous knockdown (Fig. S1, red curve) and over-expression (Fig. S1, green curve) in adults were very similar to the results described above. In summary, it appears that there is an essential requirement for Hipk in adult muscles, but not in adult dopaminergic neurons. In a pilot study, we found that Hipk over-expression in muscles of adult males did not affect survival (data not shown).

We extended the developmental analysis in a second set of experiments using a different RNAi transgene *UAS-hipk*-RNAi (V2) and a second Hipk-expressing transgene *UAS-hipk* (attP40) and a subset of nervous system and muscle *Gal4* drivers. The data indicate that, similar to the results for the NIG *hipk*-RNAi transgene, the V2 UAS-*hipk*-RNAi transgene was lethal when expressed in muscles and semi-lethal when expressed throughout the nervous system (Supplementary Table 1). However, in contrast to the previous knockdown data, V2 *UAS-hipk-RNAi*-induced knockdown in dopaminergic neurons as well as in glial cells, had no effect. In addition, in agreement with the previous over-expression data, the attP40 *UAS-hipk* transgene is lethal/semi-lethal when expressed throughout the nervous system and semi-lethal when expressed in dopaminergic neurons (Supplementary Table 2). However, in contrast to the previous overexpression data, expression of the attP40 *UAS-hipk* transgene in muscles and glial cells had no effect. Moreover, neither V2 *UAS-hipk* RNAi nor attP40 *UAS-hipk* generated a rough eye phenotype when driven with *elav-Gal4* or *longGMR-Gal4* (data not shown). We believe that the aforementioned differences in the data are due to variable potency of the V2 *UAS-hipk*-RNAi and attP40 *UAS-hipk* transgenes. Collectively, our data suggest that Hipk regulates elements of the fly nervous system and that Hipk causes lethality when mis-expressed in a subset of neurons and glial cells. In addition, the lethal effects of perturbation of Hipk in muscle indicate essential roles in the development and/or function of the tissue.

### Hipk is present in the muscle cytoplasm and nuclei during larval NMJ development, and regulates NMJ size

As modulation of Hipk levels with *Mef2-Gal4* caused pupal lethality, we examined the distribution of Hipk at the neuromuscular junction (NMJ) of 3^rd^ instar larvae. In wild-type, Hipk immunoreactivity was observed as punctate labeling throughout the muscle cytoplasm and within the DAPI-negative regions of muscle nuclei, with no obvious accumulations at the NMJ (Fig. 3A-A’”, C-C”). Muscle-specific Hipk over-expression led to increased Hipk extra-synaptic puncta (Fig 3B-B”’). Most strikingly, however, muscle nuclei displayed considerable Hipk immunoreactivity in the form of large, round bodies that consisted of strong Hipk labeling at the surface of these structures with an internal core area containing less Hipk labeling (Fig. 3B’-B’”, D-D”). These structures were present in the DAPI-negative regions of the nucleus but did not localize within the nucleolus.

**Fig. 3:**
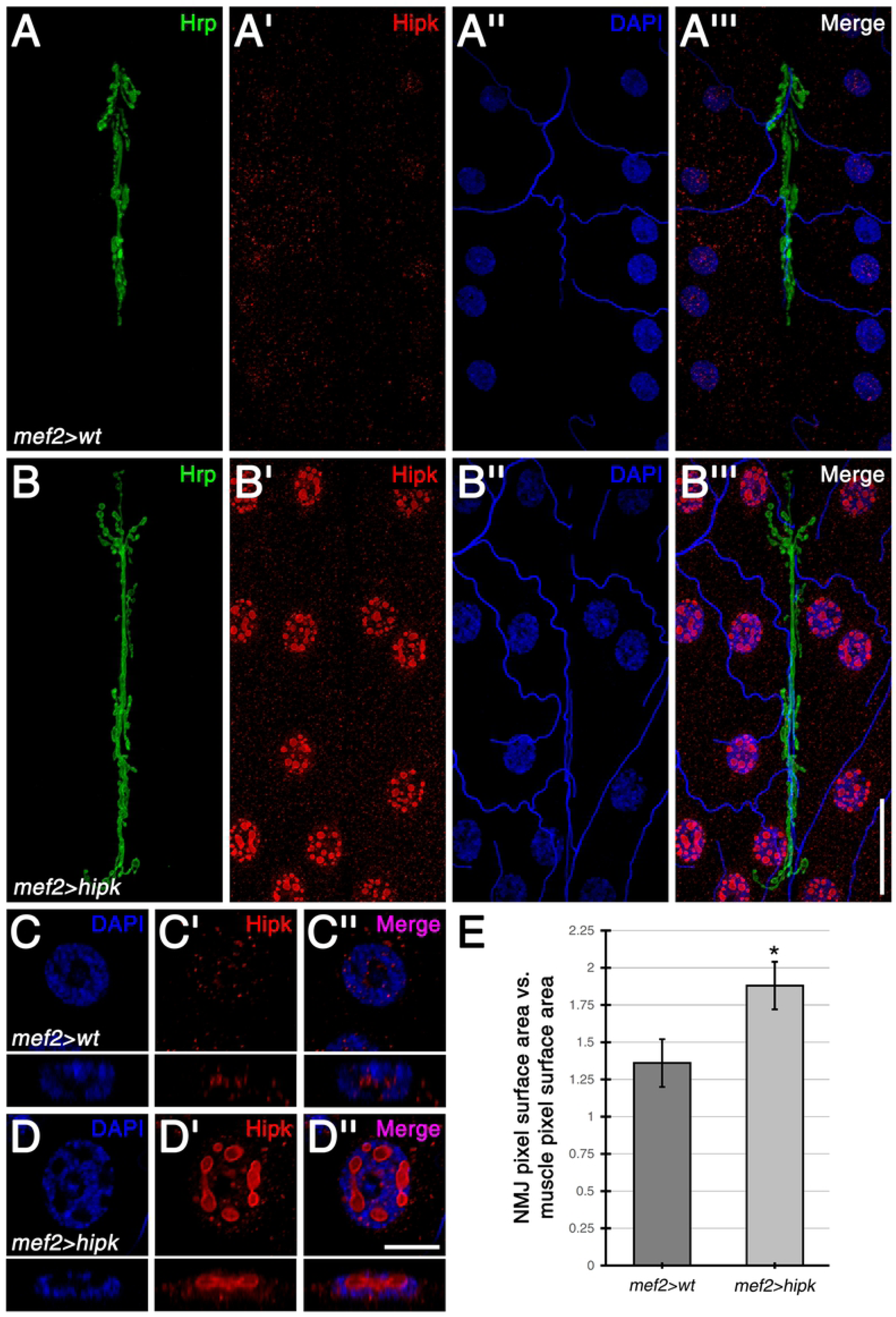
Hipk is present throughout the muscle cytoplasm and within muscle nuclei during larval NMJ development. (A-A”’) In wild-type control (i.e. *Mef2-Gal4* outcrossed to *w*^*1118*^) larval body wall muscles, Hipk showed an extra-synaptic punctate distribution throughout the muscle cytoplasm and within muscle nuclei, which were marked with DAPI. No distinct accumulations were observed at the NMJ, which was marked by Hrp. (B-B”’) Muscle-specific expression of *UAS-hipk* with *Mef2-Gal4* led to increased Hipk puncta in the muscle cytoplasm. Most strikingly, Hipk also accumulated within muscle nuclei as large, round bodies. (C-D”) Magnified, single-sectional views of muscle nuclei. For the wild-type control, Hipk puncta were found within the nucleus, primarily in DAPI-negative regions (C-C”, see cross-sectional views in the panels directly below). With Hipk over-expression in the muscle, Hipk was present at both the boundary and within the nuclear bodies, which were also localized in DAPI-negative regions (D-D”, see cross-sectional views). (E) Quantification of the effects of Hipk on NMJ size. NMJ size was measured as NMJ pixel surface area normalized against muscle pixel surface area. Over-expression of Hipk in the muscle resulted in significantly larger NMJs when compared to the wild-type control. Sample size was six NMJs per genotype. * p>0.05. Scale bars: 40µm (A-B”’); 10uM (C-D”).

Due to the presence of Hipk during larval NMJ development, we next tested for Hipk-mediated effects on NMJ size and found that the NMJs of *Mef2*-driven Hipk over-expressors were significantly larger than the NMJs of the wild-type controls (Fig 3A, B, E). Expression of transgenic *hipk-RNAi* in the muscle caused severe muscle morphological defects (Fig. S2B) and was not analyzed further in this study. As Hipk did not specifically localize to either the pre- or post-synapse of the NMJ at endogenous or over-expressed levels, Hipk regulation of NMJ size likely arises from indirect modulation of synaptic proteins.

### Hipk regulates multiple post-synaptic proteins, including the cytoskeletal protein, Hts

To elucidate Hipk’s role in regulating NMJ size in the muscle, we assessed the distribution of multiple post-synaptic proteins in Hipk over-expressing larvae. No effects on the synaptic levels of GluRIIA were observed between muscle-specific Hipk over-expression and the wild-type control (Fig. S3A). However, over-expression of Hipk resulted in significantly lower α-Spectrin levels (Fig. S3B) but higher Fasciclin 2 levels (Fig. S3C) at the NMJ. As the mode of regulation differed between the post-synaptic proteins evaluated, the observed alterations associated with Hipk over-expression appear to be specific, and not due to artificial or non-specific causes.

Another post-synaptic protein that has been shown to regulate NMJ growth is the actin-spectrin associated protein, adducin/Hts (Pielage et al., 2011; Wang et al., 2011). In wild-type control NMJs, Hts predominately localized to the postsynaptic membrane (Fig. 4A-A”). Hts was also present extra-synaptically throughout the muscle membrane as punctate structures organized in a lattice-like pattern (Fig. 4E), a distribution that is observed with spectrin networks in other systems (Han et al., 2017; Xu et al., 2013). Muscle-specific over-expression of Hipk caused no observable effects on Hts localization at the NMJ (Fig. 4B-B”). However, Hts distribution throughout the extra-synaptic muscle membrane was either undetectable or disrupted (Fig. 4F). In addition, aberrant accumulations of Hts often formed (Fig. 4G). These results indicate that Hipk negatively regulates Hts extra-synaptic distribution at the muscle membrane.

**Fig. 4:**
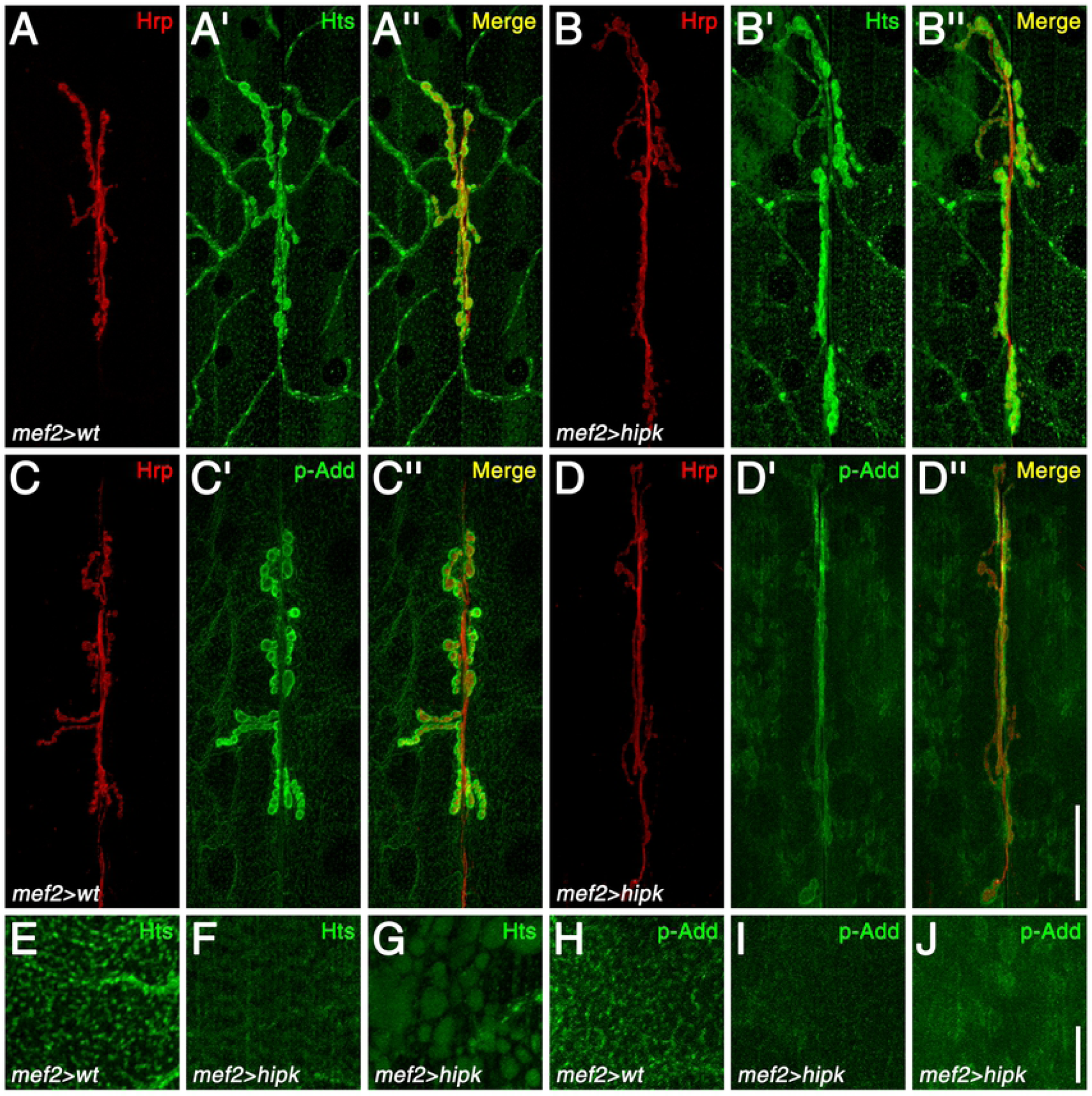
Hipk disrupts Hts distribution in the muscle and suppresses its phosphorylation at the MARCKS domain. (A-A”) In the wild-type control, Hts localized to the NMJ. In addition, Hts puncta formed a lattice-like network throughout the muscle. (B-B”) With Hipk over-expression in the muscle, Hts still localized to the post-synaptic membrane of the NMJ. However, the organization of the Hts muscle puncta was disrupted. Cloudy accumulations in the muscle were also often observed. (C-C”) p-Add immunoreactivity in the wild-type control was present at the NMJ, and was also observed as puncta that formed the lattice-like network in the muscle. The antibody detects an Add phosphorylation site in the MARCKS domain, which is conserved in Hts. (D-D”) Muscle-specific over-expression of Hipk reduced p-Add immunoreactivity, as lower signal levels were observed at the NMJ, and the lattice-like network in the muscle was missing. In addition, aberrant accumulations were observed in the cytoplasm. (E-J) High-magnification views of Hts and p-Add immunoreactivity in a section of muscle membrane, showing the lattice-like network in wild-type (E and H) and Hipk over-expression phenotypes, *i.e.* disrupted organization (F and I) and aberrant accumulations (G and J). Scale bars: 40µm (A-D”); 10µm (E-J).

In many cell types, phosphorylation of adducin/Hts by protein kinase C (PKC) or cAMP-dependent protein kinase (PKA) can inhibit the interactions of adducin/Hts with the spectrin-actin cytoskeleton, and cause adducin/Hts to redistribute from the plasma membrane to the cytoplasm (Matsuoka et al., 1998). To determine if Hipk affects Hts distribution at the muscle membrane through phosphorylation, we used an antibody that detects phosphorylation of the mammalian adducins at the PKC/PKA target site in the myristoylated alanine-rich C-kinase (MARCKS) domain. The site is conserved in *Drosophila* Hts, and the antibody has been previously used on larval muscles to detect Hts phosphorylation (Wang et al., 2014). In the wild-type control, Hts phosphorylation was observed at the NMJ, and was also present as puncta that formed the lattice-like network in the muscle described above (Fig. 4C-C”, H). Muscle-specific over-expression of Hipk suppressed Hts phosphorylation, as lower levels of p-adducin immunoreactivity were observed at the NMJ (Fig. 4D-D”), and the lattice-like network in the muscle was missing (Fig. 4I). In addition, aberrant accumulations in the muscle cytoplasm were also observed (Fig. 4J). These results indicate that Hipk influences the phosphorylation of Hts, either directly or indirectly.

### Hipk suppresses CaMKII and PAR-1 expression, thus modulating Dlg phosphorylation

We have previously shown that Hts regulates the post-synaptic localization of the important scaffolding molecule, Discs large (Dlg), by elevating the levels of the kinases, CaMKII and PAR-1, in the muscle (Wang et al., 2011). CaMKII and PAR-1 phosphorylate Dlg at the PDZ1 and GUK domains, respectively, which disrupts Dlg postsynaptic targeting (Koh et al., 1999; Zhang et al., 2007b) and influences synaptic stability (Krieger et al., 2016). To assess if Hipk modulates this function of Hts, CaMKII and PAR-1 immunoreactivity were observed in the muscles of control and Hipk over-expressing larvae. CaMKII and PAR-1 typically localize to the NMJ, but are also present throughout the muscle (Fig. 5A, C), as shown previously (Koh et al., 1999; Zhang et al., 2007b). When Hipk was over-expressed in the muscle, immunoreactivity against CaMKII and PAR-1 was reduced (Fig. 5B, D). PAR-1 immunoreactivity at the muscle membrane was missing in comparison to the wild-type control (compare corresponding cross-sections for Fig. 5D, C). CaMKII immunoreactivity at the muscle membrane was also reduced, revealing the underlying striated pattern of the sarcomeres (compare cross-sections for Fig. 5B, A). To determine if the observed decreases in CaMKII and PAR-1 immunoreactivity were due to changes at the transcript level, FISH was performed with the use of labeled antisense probes that bind to *camkII* and *par-1* mRNA. Endogenous *camkII* and *par-1* transcripts appeared as puncta throughout the muscle cytoplasm, with prominent accumulations found within muscle nuclei (Fig. 5E-E”, G-G”) (Wang et al., 2014). However, muscle-specific Hipk over-expression caused overall reductions in the levels of both transcripts (Fig. 5F-F”, H-H”; quantifications in Fig S4A, B). These results indicate that Hipk suppresses the expression of CaMKII and PAR-1 at the transcript level, thereby modifying the levels and distribution of these kinases in the muscle.

**Fig. 5:**
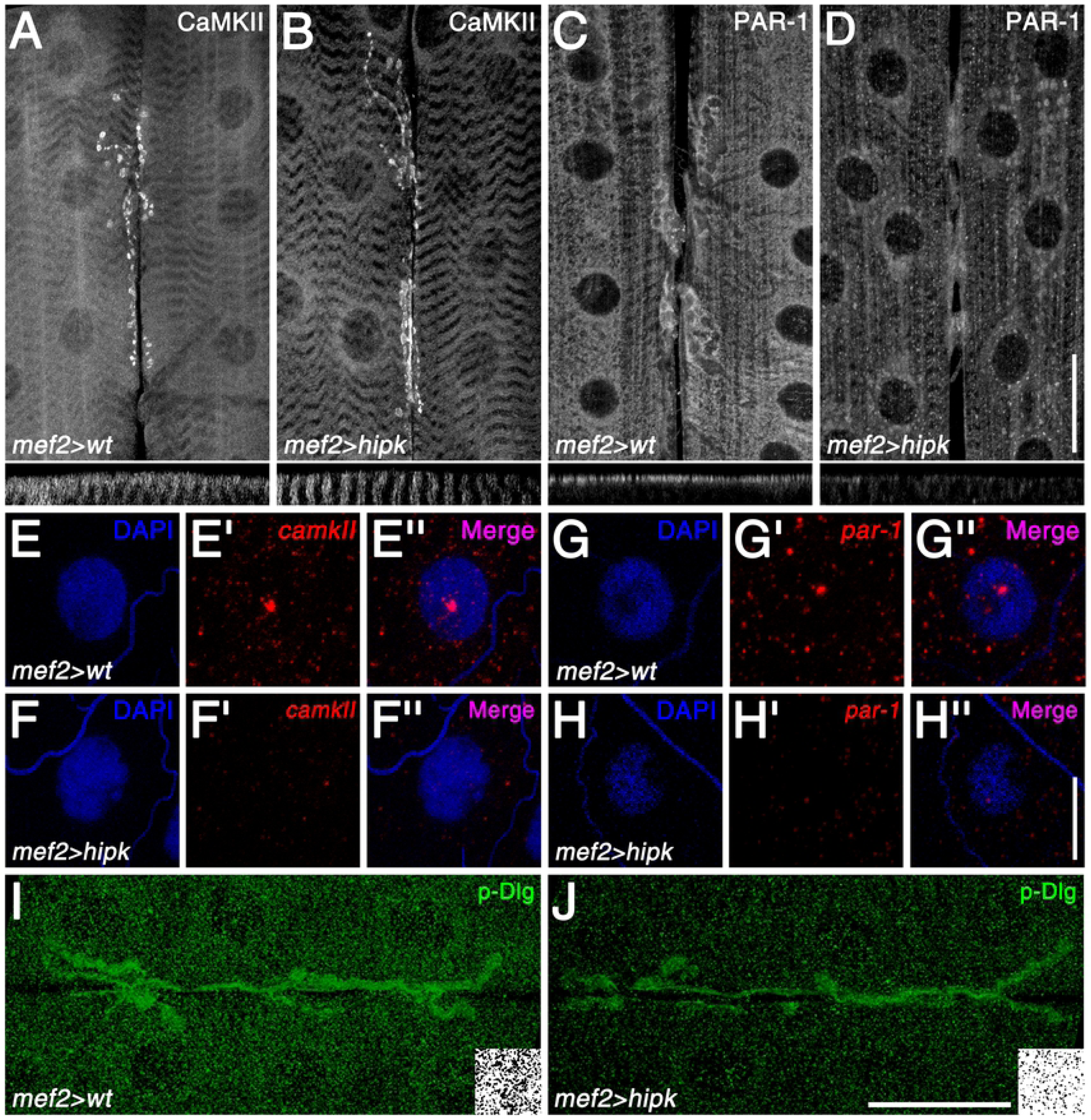
Hipk reduces Dlg phosphorylation through negative regulation of CaMKII and PAR-1 expression. (A) In the wild-type control, CaMKII localized specifically at the NMJ, but was also present throughout the muscle cytoplasmic surface (see cross-section of the muscle in the panel directly below). (B) Muscle-specific Hipk over-expression led to reduced CaMKII levels in the muscle, exposing the underlying striated pattern of the sarcomeres (more easily observed in the cross-section). (C) Similar to CaMKII, PAR-1 in the wild-type control was present at the NMJ and muscle cytoplasmic surface (see cross-section). (D) Hipk over-expression also resulted in lower PAR-1 levels in the muscle (more easily observed in the cross-section). (E-E”) In the wild-type control, *camkII* transcripts were found as distinct accumulations within muscle nuclei, with lower levels of puncta found in the muscle cytoplasm. DAPI was used to mark the nuclei. (F-F”) Over-expression of Hipk in the muscle caused a marked decrease in overall *camkII* transcript levels. (G-H”) Similar results were observed for *par-1* transcripts. (I) p-Dlg in the wild-type control was observed at the NMJ and as puncta throughout the muscle cytoplasm. The antibody detects Dlg phosphorylation at the PAR-1 target site in the GUK domain. Inset shows p-Dlg immunoreactivity in a section of muscle that was converted to a black-and-white image. (J) Hipk over-expression resulted in reduced p-Dlg puncta levels in the muscle (compare insets). Scale bars: 40µm (A-D); 20µm (E-H”); 40µm (I,J).

We next assessed whether Hipk also regulates Dlg by evaluating potential Hipk-mediated effects on Dlg phosphorylation. To accomplish this, we made use of an antibody that specifically detects Dlg phosphorylation at the PAR-1 target site (Zhang et al., 2007b). Unfortunately, no antibody against phosphorylation at the CaMKII target site was available, thus we were unable to test this directly. In the wild-type control, p-Dlg^S797^ immunoreactivity was observed at the NMJ and as puncta throughout the muscle cytoplasm (Fig. 5I), as previously reported (Wang et al., 2011; Zhang et al., 2007b). However, when Hipk was over-expressed in the muscle, extra-synaptic p-Dlg puncta levels were significantly decreased (Fig. 5J; quantification in Fig. S4C). Collectively, the data indicate that Hipk can modulate the phosphorylation of Dlg by PAR-1, and likely CaMKII as well, through negatively regulating the expression of these kinases.

### Hipk modulates the localization of nuclear proteins

The nuclear Hipk proteins have been previously associated with nuclear structures known as ‘Hipk domains’ (Moller et al., 2003), PML bodies, or paraspeckles (Rinaldo et al., 2008). To further investigate the relationship between the striking nuclear Hipk structures observed on Hipk over-expression and that of other nuclear factors, we performed immunocytochemistry with antibodies to two nuclear proteins demonstrated to have extensive roles in the nervous system. For instance, TAR DNA binding protein 43 (TDP-43) is a DNA- and RNA-binding protein predominantly localized to the nucleus that has roles in transcriptional regulation, as well as RNA processing and stability (Buratti and Baralle, 2001; Casci and Pandey, 2015). This protein is present in *Drosophila*, where it has been well characterized and is called TBPH (Casci and Pandey, 2015). Additionally, the protein fused in sarcoma (FUS/TLS), which in *Drosophila* is represented by Cabeza (Caz, or dFUS), is a nuclear-associated transcriptional activator which has also been well characterized (Lanson et al., 2011). Mutations in TDP-43 and FUS have been identified in patients with the familial form of the neurodegenerative disorder amyotrophic lateral sclerosis (ALS), which is characterized by progressive loss of motoneurons and muscle atrophy.

In the wild-type control, Caz localized predominately to muscle nuclei (Fig. 6A). Muscle-specific Hipk over-expression did not alter Caz distribution or levels (Fig. 6B). TBPH immunoreactivity in the wild-type showed a largely nuclear distribution, with some labeling of the muscle cytoplasm (Fig. 6C). In contrast, with Hipk over-expression, TBPH immunoreactivity was strikingly reduced in the nucleus, with increased TBPH labelling around the nuclear membrane (Fig. 6D). The presence of cytoplasmic TDP-43/TBPH is seen in a variety of neurological disorders including ALS, as well as fly models of ALS, and is believed to be due to the inability of TDP-43 to re-enter the nucleus (Casci and Pandey, 2015; Chou et al., 2018).

**Fig. 6:**
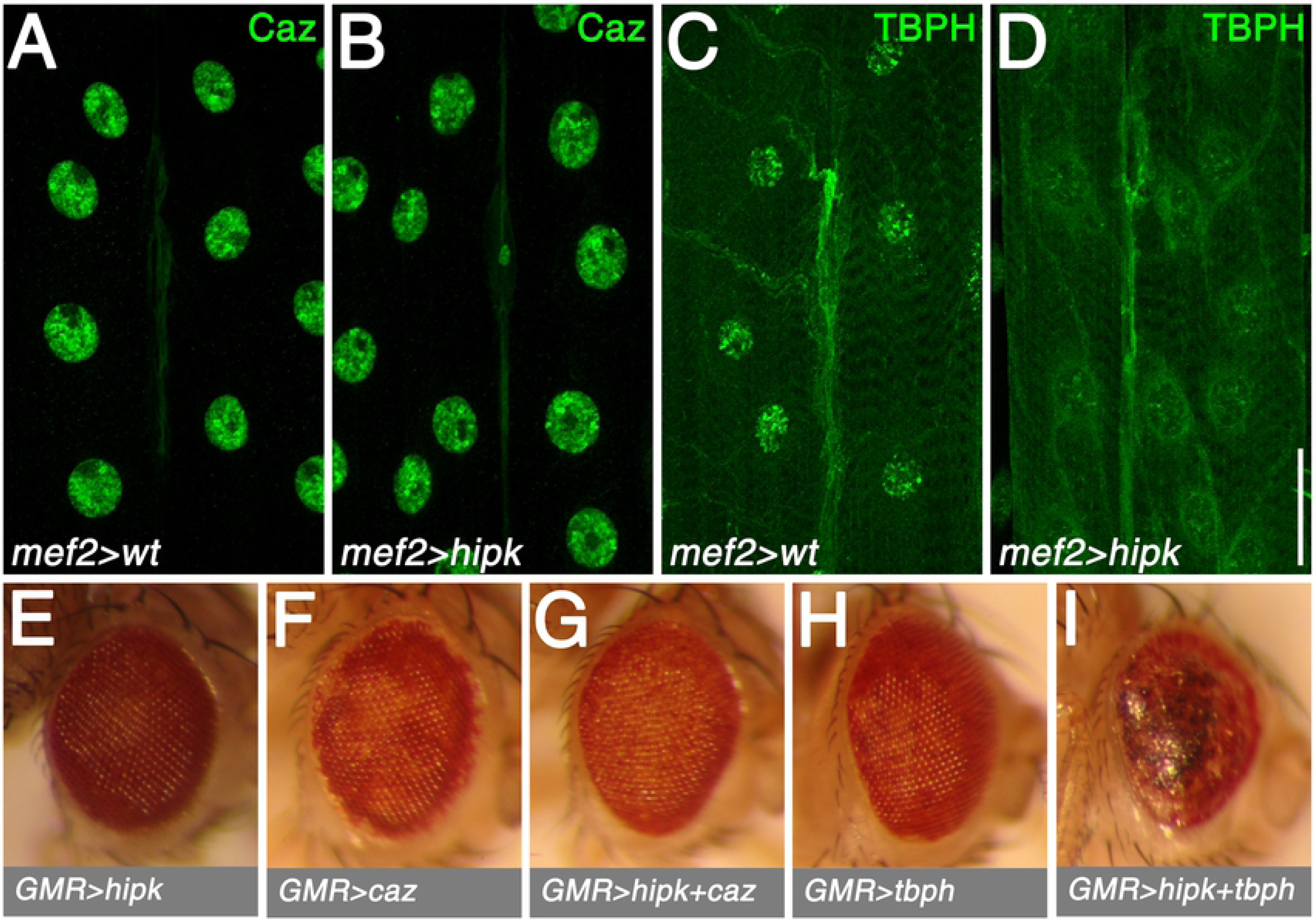
Hipk-mediated effects on the ALS-associated proteins, FUS/Caz and TDP-43/TBPH. (A) In the wild-type control, Caz localized predominately to muscle nuclei. (B) Muscle-specific over-expression of Hipk caused no observable effects on Caz distribution or levels. (C) TBPH was also most prevalent in muscle nuclei in the wild-type control. (D) Hipk over-expression in the muscle resulted in reduced nuclear TBPH levels. (E-I) Genotypes for eye images (all males incubated at 29°C): *UAS-hipk/*+; *longGMR-Gal4/*+ (E), *UAS-caz/CyO; longGMR-Gal4/*+ (F), *UAS-hipk/UAS-caz; longGMR-Gal4/*+ (G); *CyO/UAS-TBPH; longGMR-Gal4/*+ (H), and *UAS-hipk/UAS-TBPH; longGMR-Gal4/*+ (I). Scale bar: 40µm (A-D).

A deleterious interaction between Hipk overexpression and TBPH overexpression was seen when Hipk (using attP40 *UAS-hipk*, which itself does not cause an eye phenotype when driven with *longGMR-Gal4*) was co-expressed with *UAS-TBPH* in *Drosophila* eyes, using *longGMR-Gal4*, leading to a rough/necrotic eye phenotype (Fig 6I). On the other hand, analogous co-expression of *UAS-caz* had no striking effect (Fig 6G).

## Discussion

Previous work has shown that elevated expression of Hipk leads to tumorigenesis and metastatic cell behavior, likely by stimulating various signaling pathways such as Wnt/Wingless, Hippo, Notch and JNK (Blaquiere et al., 2018; Chen and Verheyen, 2012; Huang et al., 2011; Lee et al., 2009a, 2009c; Poon et al., 2012). However, relatively little is known regarding the action of Hipk in the nervous system and muscle. We find that ubiquitous over-expression and knockdown of Hipk have dramatic effects on adult survival, leading to progressive immobility and death and these effects are most likely due to perturbations in the nervous system and muscle. Evidence for nervous system involvement during development is supported by the observation of rough eye phenotypes associated with pan-neural and/or eye-specific Hipk knockdown and over-expression, as well as by the fact that such mis-expression in dopaminergic neurons and glial cells is lethal. Taken together, these results show that both reduced and excessive levels of Hipk have profound effects on nervous system function in flies. Our adult-specific data are consistent with observations that *hipk2* knockout mice exhibit increased perinatal lethality and that a mutation in the *C. elegans* ortholog, *hpk-1*, shortens lifespan (Berber et al., 2016; Chalazonitis et al., 2011). The effects of Hipk misexpression in dopaminergic neurons is of significant interest since these neurons are affected by human neurodegenerative disorders such as Parkinson’s disease. Such effects are consistent with the previously mentioned role of mammalian Hipk2 in survival of midbrain dopaminergic neurons (Zhang et al., 2007a).

### Effects of Hipk on muscle

Our results demonstrate that Hipk is present in muscle nuclei where Hipk likely phosphorylates transcription factors and other substrates. Interestingly, we have also found that muscle-specific knockdown in *Drosophila* adults dramatically shortens their lifespan and that death is preceded by progressive immobility supporting our claim that muscle and neuronal function are regulated by Hipk. Under control conditions Hipk is predominantly intranuclear where it is found in small puncta, as described previously (Huang et al., 2011). As previous work did not clearly localize these puncta to identified subnuclear structures they have been referred to as ‘Hipk domains’ (Moller et al., 2003). In some cases, Hipk localization has been associated with the subnuclear promyelocytic leukemia (PML) body where Hipk2-mediated phosphorylation of p53 occurs (Moller et al., 2003). The exact nature of the Hipk nuclear domains is still unresolved, and the appearance of the nuclear structures differs between endogenous proteins and following ectopic expression of Hipk. In *Drosophila*, nuclear localization of Hipk is dependent on the small ubiquitin-related modifier (SUMO) protein (Smt3) and the sumoylation-controlled balance between nuclear and cytoplasmic Hipk has been shown to regulate the JNK signaling pathway (Huang et al., 2011). We found that overexpression of Hipk in muscle results in very large nuclear Hipk-domains having a ‘rim’ of strong Hipk immunoreactivity with less labelling in the core of these structures. As these structures were much larger than the Hipk domains in wild-type, it is possible that they comprise regions of aggregated Hipk. The structures were confined to the nucleus, with no evidence of large cytoplasmic Hipk-positive structures. Within the nucleus the Hipk domains were extra-nucleolar and were exclusively present in the extrachromosomal regions of the nucleus.

### Effects of Hipk on NMJ

To evaluate the possible mechanism of Hipk action at the NMJ and muscle we studied NMJs in Hipk over-expressing animals and found that terminal axonal branching onto these muscles was significantly larger than controls. We found that these NMJ boutons had a striking reduction in levels of phosphorylated Hts, which could be related to the axonal branching effect. Adducin/Hts is an actin- and Spectrin-binding protein that is present both pre- and post-synaptically in the fly NMJ. It is regulated by phosphorylation and we have found that phosphorylated Adducin (p-Add) modulates levels of transcripts of the important protein kinases CaMKII and PAR-1 (Wang et al., 2011, 2014). Thus, our observations that Hipk is associated with reductions in p-Add at the synapse as well as decreased transcripts and immunoreactivity of CaMKII and PAR-1 in the synapse might be of functional significance and a consequence of Hipk action. We would predict that the reductions in phosphorylation of Adducin with Hipk overexpression cause Add to remain at the synapse and lead to the increase in NMJ surface area observed in the Hipk over-expressors.

To further extend observations on the Hipk nuclear functions we examined several proteins that are well known to be expressed in fly muscle nuclei including Caz and TBPH/TDP-43. These proteins have been studied extensively, as TDP-43 redistribution from the nucleus to the cytoplasm is frequently observed both in the neurodegenerative disorder, amyotrophic lateral sclerosis (ALS) and in fly models of ALS (Casci and Pandey, 2015; Chang and Morton, 2017). In wild-type tissue, TBPH is enriched in nuclear structures or aggregates, some of which overlap with endogenous Hipk proteins. We evaluated TBPH localization in individual Hipk over-expressing larval muscle nuclei. Under these conditions, we observed a striking decrease in nuclear TBPH immunoreactivity and de-localization of TBPH such that the nuclear and cytoplasmic levels were similarly low. These findings suggest that Hipk may regulate TBPH distribution either directly or indirectly. In contrast, another nuclear protein involved in neurodegeneration, Caz, is not clearly affected by Hipk modulation suggesting that the effects we observe with TBPH are not simply generic effects on nuclear proteins.

Our results are also of interest in relation to recent work showing that activation of Hipk2 promotes endoplasmic reticulum (ER) stress and neurodegeneration in an animal model of ALS (Lee et al., 2016). The effect of Hipk2 in ALS may be mediated by TDP-43, as well as the JNK signaling pathway. Cytosolic TDP-43 is often found to be phosphorylated in tissue from ALS patients and an elevation of phosphorylated Hipk2 (at S359/T360) was found to be positively correlated with the level of ubiquitinated TDP-43 in tissue from ALS patients, post-mortem (Lee et al., 2016). Hipk2 inhibition reduced the neurodegeneration associated with ER stress *in vitro* produced either by tunicamycin or by altered expression of wt or mutant TDP-43 (Lee et al., 2016). Interestingly, mice having mutations in FUS/TLS that are associated with familial ALS did not demonstrate the altered Hipk2 levels, consistent with our observations that Hipk over-expression did not alter the distribution of nuclear Caz [*Drosophila* FUS], or produce a rough eye phenotype with Caz over-expression. However, it is possible that given the many signaling pathways that are triggered by Hipk, other potential mechanisms for neurotoxicity exist. For instance, Hipk2 has also been linked to transcriptional control of N-methyl-D-aspartate receptors, which are well established to be involved in neuronal cell death (Shang et al., 2018).

Our data demonstrate that in addition to having roles in tumorigenesis Hipk also has profound effects on the nervous system and muscle including on the NMJ. This work also indicates that in addition to modifying phosphorylation of nuclear proteins, Hipk affects the distribution of nuclear proteins and also can influence the phosphorylation state of extra-nuclear proteins, either directly or indirectly affecting function and viability.

## Materials and Methods

### Fly Stocks

All stocks were maintained at 25°C or 18°C. The *UAS-hipk*-RNAi (NIG) line (17090R-1) used for most of the current work was obtained from the NIG stock centre. A second *UAS-hipk*-RNAi (V2) line (108254) used for some crosses was obtained from the VDRC stock centre. *UAS-HA-hipk* is described in (Lee et al., 2009a) and attP40 *UAS-HA-hipk* in Tettweiler et al., 2019 (in preparation). The following lines were obtained from the Bloomington *Drosophila* Stock Center: *w*^*1118*^ (3605); *w*^1118^; *P{UAS-Dcr-2.D}2* (*Dicer-2* on chromosome 2; 24650); *w*^*^; *P{tubP-Gal80*^ts^}*20*; *TM2*/*TM6B, Tb* (*Gal80*^*ts*^ on chromosome 3; 7019 *P{tubP-Gal80*^ts^}*2* (*Gal80*^*ts*^ on chromosome 3; 7017); *y w*; *P{Gal4-Mef2.R}3 (Mef2-Gal4*; 27390); *ple-Gal4* (8848); *ChAT-Gal4.7.4-19B* (6798); *ChAT-DD. Gal4.VP16* (64410); *GawB elav*^C155^ (*elav-Gal4*; 458); *GawB V Glut*^OK371^ (26160); *GawB D42* (8816); *repo-Gal4* (7415); *w*^*^; *P{longGMR-Gal4}2* (*longGMR-Gal4*; *8605*); *y w*; *wg*^Sp-1^/*CyO*; *P{longGMR-Gal4*}*3*/*TM2* (*longGMR-Gal4*; 8121). The *Appl-Gal4* line was kindly provided by Dr. U. Pandey. The *UAS-TBPH* and *UAS-caz* lines were kindly provided by Dr. B. McCabe. The *TH-Gal4* line was kindly provided by Dr. D. Allan. The following stocks were constructed for the adult survival experiments and analysis of eye effects: (i) *UAS-hipk*-RNAi/UAS-*hipk*-RNAi; *Gal80*^*ts*^/*Gal80*^*ts*^ (ii) *Gal80*^*ts*^/*Gal80*^*ts*^; *UAS-HA-hipk*/*UAS-HA-hipk* (iii) *Gal80*^*ts*^/*Gal80*^*ts*^; *Tub-Gal4*/*TM3, Sb* (iv) *Dicer-2/Dicer-2*; *Tub-Gal4/TM3, Sb Ser*; (v) attP40 *UAS-HA-hipk*/*CyO; longGMR-Gal4*/*longGMR-Gal4*.

### Fly crosses, lifespan and phenotypic analyses

Two different sets of crosses were used for lifespan analysis. In the first set, homozygous UAS*-hipk-RNAi* (NIG) or *UAS-HA-hipk* females were crossed to *Gal80*^*ts*^/*Gal80*^*ts*^; *Tub-Gal4*/*TM3, Sb* males. In the second set, doubly homozygous *UAS-hipk-RNAi* (NIG); *Gal80ts* males were crossed separately to *Dicer-2*/*Dicer-2*; *Tub-Gal4*/*TM3, Sb* females or homozygous *Mef2-Gal4* or *TH-Gal4* females. These crosses were set up at 18°C and the parents were sub-cultured twice or three times to fresh medium at three-day intervals and the progeny were allowed to develop to eclosion at 18°C. Relevant experimental and internal control males from these crosses (see the Results), as well as Oregon-R (+) control males raised under the same conditions, were collected within 0-24 hours post-eclosion and shifted to 29°C (10 flies/vial). Thereafter, the males were transferred to fresh medium, usually every 2 or 3 days, and the number of survivors was recorded at each transfer, in most cases until the last of the flies had died. The data were converted to percent survival and these values were plotted versus days at 29°C. For the first set of crosses, there were two trials each for knockdown, over-expression and two types of internal controls and three trials for Oregon-R controls. Lethal phase (LP) refers to the particular stage of development (e.g. larvae, pupae) when death usually occurred.

Crosses were conducted to test for possible developmental effects of Hipk knockdown or Hipk over-expression in the nervous system and muscles at 29°C in two separate sets of experiments. In the first, most crosses involved mating *Gal4*-driver-bearing males to heterozygous females bearing either NIG *UAS-hipk-RNAi* or *UAS-hipk* transgenes and appropriate balancer chromosomes (*CyO* or *TM3, Sb* or *TM3, Sb Ser*); the single exception involved the glial cell crosses, which used repo-*Gal4*/*TM3, Sb* females and *hipk*-transgene-bearing males. In the second set of experiments, most crosses involved mating females bearing either V2 *UAS-hipk*-RNAi/CyO or attP40 *UAS-hipk* /CyO to *Gal4*-driver-bearing males; the single exception involved tests with the *Appl-Gal4* driver, for which reciprocal crosses were performed. In all cases, crosses were carried out in vials with ∼four pairs of parents per vial and the crosses were sub-cultured on fresh medium twice at 3-day intervals. Wherever crosses showed lethality, approximate lethal phases were determined, and any phenotypes of survivors were also noted. In addition, with respect to the first set of experiments, the eyes of surviving *elav-Gal4*/+; *UAS-hipk-RNAi*/+, *elav-Gal4*/+; *UAS-hipk*/+ females, as well as those of respective internal control *elav-Gal4*/+; *CyO*/+ and *elav-Gal4*/+; *TM3, Sb*/+ flies, were photographed.

To test for eye-specific effects of Hipk knockdown and over-expression, NIG *UAS-hipk*-RNAi/*CyO* and 3M *UAS-hipk*/*TM3, Sb* females were crossed separately to *longGMR-Gal4* (chromosome 2) males at 29°C (four pairs of parents per vial) and the crosses were sub-cultured on fresh medium twice at three-day intervals. *UAS-hipk-RNAi*/*longGMR-Gal4* flies failed to eclose at 29°C (data not shown; see Results). After eclosion, the eyes of *longGMR-Gal4/*+; *UAS-hipk*/+ females and of internal control *CyO*/*longGMR-Gal4* and *longGMR-Gal4*/+; *TM3, Sb*/+ females, were photographed. Since pilot studies revealed that attP40 *UAS-hipk*/*longGMR-Gal4 f*lies developing at 29°C did not exhibit an eye phenotype (data not shown), we used this line to test for potential interactions involving simultaneous Hipk over-expression and over-expression of nuclear proteins of interest, Caz and TBPH. For this analysis, attP40 *UAS-HA-hipk*/*CyO*; *longGMR-Gal4*/*longGMR-Gal4* females were mated separately to homozygous *UAS-caz* and *UAS-TBPH* males at 29°C (four pairs of parents per vial) and the crosses were sub-cultured on fresh medium twice at three-day intervals. In all cases, after eclosion the eyes of appropriate experimental and internal control flies were photographed (see Results for genotypes). In addition, the eyes of surviving *elav-Gal4*/+; *UAS-hipk-RNAi*/+ and *elav-Gal4*/+; *UAS-hipk*/+ and appropriate internal control females from the aforementioned developmental crosses, were also photographed.

### Larval Body Wall Preparation

Body wall dissections of crawling third instar larvae were performed as previously described (Wang et al., 2015). For transgenic analysis, homozygous *UAS*-transgene-bearing males were crossed to homozygous *Gal4*-bearing virgin females ensuring that all progeny carried one copy of each. All crosses were maintained at 25°C unless indicated otherwise.

### Immunohistochemistry (IHC)

Immunostaining of body walls was performed as previously described (Ramachandran and Budnik, 2010). The following primary antibodies were used: 1:100 goat anti-Hrp (Jackson ImmunoResearch – 123-005-021); 1:100 rabbit anti-Hrp (JIR – 323-065-021); 1:200 rabbit anti-Hipk (Blaquiere et al., 2018); 1:50 mouse anti-GluRIIA (Developmental Studies Hybridoma Bank – 8B4D2); 1:10 mouse anti-α-Spec (DSHB – 3A9); 1:2 mouse anti-Fas2 (DSHB – 1D4); 1:10 mouse anti-Hts (DSHB – 1B1); 1:100 goat anti-phospho-Add^S662^ (Santa Cruz Biotechnology – sc-12614); 1:200 mouse anti-CaMKII (Cosmo Bio – CAC-TNL-001-CAM); 1:200 rabbit anti-PAR-1(Sun et al., 2001), which was generously provided by Dr. Bingwei Lu (Stanford University); 1:200 rabbit anti-phospho-Dlg^S797^ (Zhang et al., 2007b), also from Dr. Lu; 1:200 rabbit anti-Caz (Sasayama et al., 2012), kindly given by Dr. Takahiko Tokuda (Kyoto Prefectural University); and 1:200 guinea pig anti-TBPH (Swain et al., 2016), a gift from Dr. Dale Dorsett (Saint Louis University). Fluorescent-labeled secondary antibodies from JIR and Vector Laboratories were all used at a 1:200 dilution. Stained body walls were mounted in VECTASHIELD Mounting Medium with DAPI (Vector – H-1200). Experiments and their controls were immunostained in the same tube, and thus exposed to the same treatment. Images of NMJs innervating muscles 6/7 from abdominal segment 3 were taken as merged stacks, unless otherwise stated, on either a Nikon A1R laser scanning confocal microscope with NIS-Elements software or a Zeiss LSM 880 with Airyscan, with experiments and their controls imaged under identical acquisition settings. All images were processed with Adobe Photoshop.

### Fluorescent in situ Hybridization (FISH)

FISH of body walls was performed as previously described (Wang et al., 2014). cDNA clones (*Drosophila* Genomics Resource Center) IP15240 and RE47050 were used as templates to make DIG-labeled, antisense probes against *camkII* and *par-1* transcripts, respectively. Both cDNA clones were obtained from the *Drosophila* Genomics Resource Center. Experiments and their controls were processed in the same tube, and thus exposed to the same treatment. Body walls were mounted and imaged as described in the previous section.

### Quantifications

Stacked confocal images of larval muscles 6/7 from abdominal segment 3 were used for measurements. Student’s t-test was used for statistical analysis, and data was expressed as the mean ± S.E.M. <u>NMJ size</u>: For each NMJ, Hrp immunofluorescence signal was delineated using Photoshop tools and the size was measured as pixel surface area. The outline of muscles 6/7 was also selected with Photoshop and the pixel surface area was recorded. To normalize the NMJ size measurement, the NMJ pixel surface area was divided by the muscle pixel surface area. <u>Immunofluorescence intensity at the NMJ</u>: Quantification of post-synaptic protein levels at the NMJ was calculated as a ratio between the fluorescence intensity of the protein of interest and Hrp, which was used as a control. Immunofluorescence signal at the NMJ was selected using Photoshop, and the intensity was determined by measuring the mean gray value. <u>Puncta levels in the muscle</u>: For quantification of *camkII*/*par-1* FISH puncta or p-Dlg IHC puncta in the muscle, confocal images were inverted and the threshold function of ImageJ was used to create binary images in which puncta were represented by black pixels and the background was eliminated. The number of extra-synaptic black pixels within a fixed area size was measured in both muscles 6 and 7.

## Acknowledgments

We are grateful to the Bloomington *Drosophila* Stock Center (NIH P40OD018537), the Vienna *Drosophila* RNAi Center (VDRC), National Institute of Genetics (NIG), *Drosophila* Genomics Resource Center and Developmental Studies Hybridoma Bank for providing fly strains, cDNA clones and antibodies. We thank Dr. Leslie Griffith (Brandeis University) for her correspondence regarding CaMKII antibodies. This work was funded by a Discovery Grant from the Natural Sciences and Engineering Research Council of Canada (NSERC) to E.M.V., to N.H., and by grants to C.K. from NSERC and from Amyotrophic Lateral Sclerosis Canada (ALS Canada)/ Brain Canada.

---

**Fig. S1: Knockdown of Hipk in adult muscles shortens lifespan, but knockdown in adult dopaminergic neurons has no effect.** Adult males subjected to muscle specific Hipk knockdown at 29°C (*UAS-hipk-RNAi*; *Mef2-Gal4*: NIG *UAS-hipk-RNAi*/+; *Mef2-Gal4*/*Gal80*^ts^; purple curve; n= 110) died within 11-15 days. Adult Oregon R control males (black curve; n= 110) under identical conditions died much later. Adult males subjected to dopaminergic knockdown at 29°C (*UAS-hipk-RNAi*; *TH-Gal4*: *UAS-hipk-RNAi*/+; *TH-Gal4*/*Gal80*^ts^; yellow curve; n= 109) showed a lifespan that was similar to that of control males (black curve). The shortened lifespans of males subjected to Hipk knockdown (*UAS-hipk-RNAi*: *UAS-hipk-RNAi*/*Dicer-2*; *Tub-Gal4*/*Gal80*^ts^; red curve; n= 59) or over-expression (*Dicer-2*/*Gal80*^ts^; *UAS-hipk*^+^/*Tub-Gal4*; green curve; n= 110) at 29°C were comparable to those described above.

**Fig. S2: Hipk knock-down in the muscle during larval NMJ development.** (A) A typical 3^rd^ instar larval NMJ, marked by Hrp, innervating muscles 6/7 in abdominal segment 3. (B) Muscle-specific expression of transgenic *hipk-RNAi* resulted in morphological defects in the muscle. Scale bar: 40µm (A,B).

**Fig. S3: Hipk-mediated effects on different post-synaptic proteins.** Post-synaptic protein levels at the NMJ was calculated as a ratio between the fluorescence intensity of the protein of interest and Hrp, which was used as a staining control. Immunofluorescence signal at the NMJ was selected using Photoshop, and the intensity was determined by measuring the mean gray value. (A) No significant effects on the levels of GluRIIA at the NMJ were observed between muscle-specific Hipk overexpression and the wild-type control. n = 8 (over 4 body walls) for each genotype. (B) However, Hipk over-expression resulted in significantly lower synaptic levels of α-Spec. n = 6 (over 3 body walls) for each genotype. (D) Hipk over-expression also resulted in significantly higher Fas2 synaptic levels. n = 6 (over 3 body walls) for each genotype. *** p < 0.0001.

**Fig. S4: Quantification of FISH and IHC puncta levels in the muscle.** Confocal images were converted to binary images, where immunofluorescent puncta were represented as black pixels and the background was represented as white pixels (examples are shown). The number of extra-synaptic black pixels within a fixed area size was measured. (A) Muscle-specific Hipk over-expression led to significantly reduced levels of *camkII* FISH signal when compared to the wild-type control. n = 20 (over 5 body walls) for each genotype. (B) Similar results were observed for *par-1* FISH signal. n = 16 (over 4 body walls) for each genotype. (C) p-Dlg IHC signal in the muscle was also significantly lowered when Hipk was over-expressed in the muscle. n = 16 (over 4 body walls) for each genotype. * p = 0.019; ** p = 0.0007; *** p < 0.0001. Scale bars: 40µm (A); 40µm (B); 40µm (C).

